# Comparison of decidual vasculopathy in central and peripheral regions of placenta with implication of lateral growth and spiral artery remodeling

**DOI:** 10.1101/2020.06.16.154484

**Authors:** Peilin Zhang

## Abstract

**Background:** Decidual vasculopathy at late gestation was shown to be associated with spiral artery remodeling at implantation. How placental lateral growth is related to spiral artery remodeling in spatiotemporal fashion is to be investigated.

**Design:** The central and peripheral portions of 105 placentas with decidual vasculopathy at term were examined with or without preeclampsia to see if temporal vascular regeneration was present. Central and peripheral vasculopathy and central and peripheral regeneration were compared.

**Result:** Peripheral portion showed more decidual vasculopathy (88 of total 105, 83.8%) than central portion (72 of total 105, 68.6%, p=0.0018). However, central portion showed more vascular regeneration (51 of total 105, 48.6%) than peripheral portion (23 of total 105, 21.9%, p=0.0024). There is no difference in vasculopathy or regeneration with or without preeclampsia.

**Conclusion:** Spiral artery remodeling is non-synchronous during lateral growth and vascular regeneration. This spatiotemporal sequence may help interpretation of morphologic changes of decidual vasculopathy.

## Introduction

Placental growth is continuous throughout the pregnancy to reach maturity at term [1, 2]. The size and weight of the placenta are independent factors affecting the fetal growth and fetal wellbeing [3-7]. Placenta in a large part demonstrates a lateral growth pattern due in part to the mechanical limit of uterine cavity with continuous expansion of a number of cotyledons from the periphery [1]. Like many other mammalian organs, there are regulatory mechanisms of size control and cell number control in the growth of placenta [8-10]. Development of cotyledons laterally from the center involves development of new anchoring villi, trophoblastic cell shell and column with corresponding spiral artery remodeling so that the fetal villous vessels are connected to the uteroplacental intervillous circulation [1]. It is reasonable to believe that there is a time sequence (temporal sequence) in regard to spiral artery remodeling from the center where spiral artery remodeling occurs first and peripheral part of the placenta later (spatial sequence), although the spatiotemporal sequence of placental development is not frequently studied [11, 12]. Spiral artery remodeling is a significant vascular transformation of maternal vessels to accommodate the embryonic and placental growth and to establish the utero-placental circulation [13]. Spiral artery remodeling is a temporal but dynamic development process predominantly occurring in the first and second trimester [1]. Based on the previous studies there is no morphologic evidence of spiral artery remodeling after the end of second trimester [14]. The size of the placenta increases to about 20 cm in diameter and 25 - 40 cotyledons at term from a blastocyst at the beginning of implantation, although the most significant placenta weight increase occurs in the third trimester [1]. The questions remain if the spiral artery in the central portion of the placenta and those in the peripheral portion show synchronous morphologic changes of remodeling, and how this difference of spiral artery remodeling in time between the central and peripheral portions of placenta affects the development of decidual vasculopathy at late gestation, and how this spatiotemporal spiral artery remodeling affects the interpretation of morphologic changes of placenta in normal pregnancy and complications. The spatiotemporal difference is typically not an issue for most placental pathologists when the placental surface is examined, and decidual vasculopathy found as the entire placenta can be easily accessed after delivery. However, the placental bed biopsy interpretation of maternal vessels will be affected significantly due to the tissue sections obtained from the biopsy sites at the placental bed [15]. The central portion of the placental bed is emphasized for placental bed biopsy in order to achieve the expected results [15]. Under these circumstances, spatiotemporal sequence of maternal vascular changes makes difference in interpretation of the material obtained for the procedure [15]. Furthermore, the theory of “failure to invade” has been proposed as the key pathogenic mechanism of preeclampsia and hypertensive disorders of pregnancy, and the cardinal vascular change of “failure to invade” theory is the lack of trophoblastic invasion of vascular wall in the superficial myometrium. In this setting, an attempt is made to delineate the difference of morphologic features of the maternal vessels within the center and the peripheral areas of maternal surface in regards to the presence and absence of decidual vasculopathy and vascular regeneration/restoration. A slight modification of standard placental examination protocol for a period of this study was made in order to answer this question.

## Material and methods

The study is exempt from Institutional Review Board (IRB) approval according to section 46.101(b) of 45CFR 46 which states that research involving the study of existing pathological and diagnostic specimens in such a manner that subjects cannot be identified is exempt from the Department of Health and Human Services Protection of Human Research Subjects. Placentas submitted for pathology examination for a variety of clinical indications were included in the study and examined under the standard protocol with the following modification of gross dissection and examination. Standard four sections were submitted for microscopic examination with one maternal surface section at the center and one at the periphery using the horizontal dissection method previously described (Figure 1) [16]. The total of 4 sections per placenta was included with one full thickness section from the peripheral area excluding the margins and one section of umbilical cord and membrane roll (Figure 1). Routine examination of placenta was performed under the clinical practice guideline using the Amsterdam criteria [17]. Diagnostic criteria for decidual vasculopathy was according to the guideline previously described [17, 18]. Vascular regeneration / restoration (recovery) of maternal spiral artery is defined by the presence of endothelium and smooth muscle within the vascular lumen as well as intramural or endovascular trophoblasts. The presence of intramural and/or endovascular trophoblasts (acute atherosis) and fibrinoid medial necrosis of vascular wall were interpreted as the presence of decidual vasculopathy [1, 18, 19]. Immunohistochemical staining procedures were performed on paraffin-embedded tissues using Leica Biosystems Bond III automated immunostaining system following the manufacturer’s instructions. Monoclonal antibodies against CD56, WT1, smooth muscle myosin heavy chain (SMYOHC) were purchased from Agilent (Dako Corporation) for clinical in vitro diagnostics with appropriate dilutions and controls under catalogue numbers M730401-2, M3561, and M7165. Selective cases of decidual vasculopathy were confirmed by the presence of CD56 for endovascular or intramural trophoblasts, and the regeneration by presence of WT1 and smooth muscle myosin heavy chain (SMYOHC), as WT1 gene expression is known to be present in endothelial cells and smooth muscle cells of vessels [20, 21]. Immunostaining for these markers was only used in selected cases to illustrate the usefulness of the techniques, and the data was based on the morphologic features.

**Figure 1.**
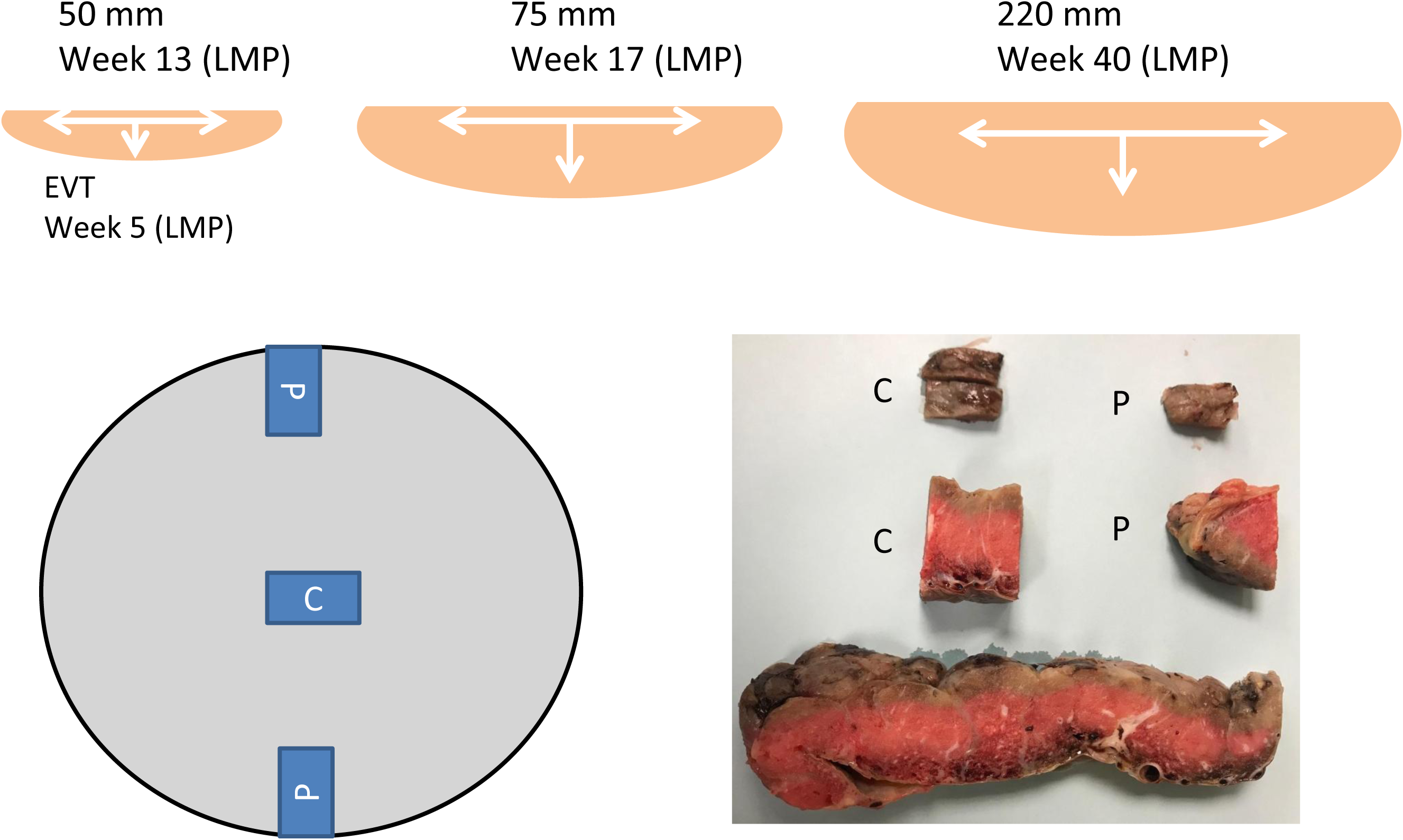
Schematic representation of placental lateral growth from early pregnancy to term with representative dissection techniques of Khong for placental surface and decidual vessels. C – central region, P – peripheral region of maternal surface

## Result

The diagram (Figure 1) showed the theoretical placental lateral growth with expansion from the center to the periphery. Endovascular trophoblasts (EVT) can be identified within the spiral artery at the week 5 from the last menstrual period (LMP). The bottom panel showed the method of gross dissection of placental surface at the center and the periphery using the method of Khong for better sampling and visualization of decidual vessels [16]. Using this method of gross sectioning, we have examined 105 placentas including 14 placentas from preeclampsia and 91 placentas from other complications without preeclampsia. The baseline characteristics of all the placentas with related clinical and pathologic findings were shown in Table 1. The baseline characteristic data from the current study was similar to those reported previously [18]. The presence of decidual vasculopathy at the center and the periphery of placentas can be easily identified under light microscope (Figure 2). The endovascular and intramural trophoblasts associated with decidual vasculopathy can be identified by using immunostaining of CD56 (Figure 3). The endothelial and smooth muscle cell regeneration / restoration within the lumen of spiral artery were detected by using WT1 and smooth muscle myosin heavy chain (SMYOHC). Co-existence of endovascular trophoblasts and endothelial cells was seen occasionally within the same spiral artery. Intramural trophoblasts with overlying endothelial cells were more frequently identified (Figure 3). There were 72 placentas with decidual vasculopathy in the central region including 59 cases with classic type decidual vasculopathy (atherosis and fibrinoid medial necrosis) and 13 cases with mixed type decidual vasculopathy (classic type and arterial mural hypertrophy) (Table 2). Totally 88 placentas in the peripheral regions showed the presence of decidual vasculopathy including 73 placentas with classic type and 15 placentas with mixed type vasculopathy (Table 2). Frequently both the central and peripheral regions of the placentas showed the presence of decidual vasculopathy but the vascular recovery (regeneration/restoration) was identified at much lower frequency at the peripheral region than in the central region (Table 1 and Table2)(p=0.0018 and 0.0024, Fisher exact probability test, one tailed and two tailed).

**Table 1:**
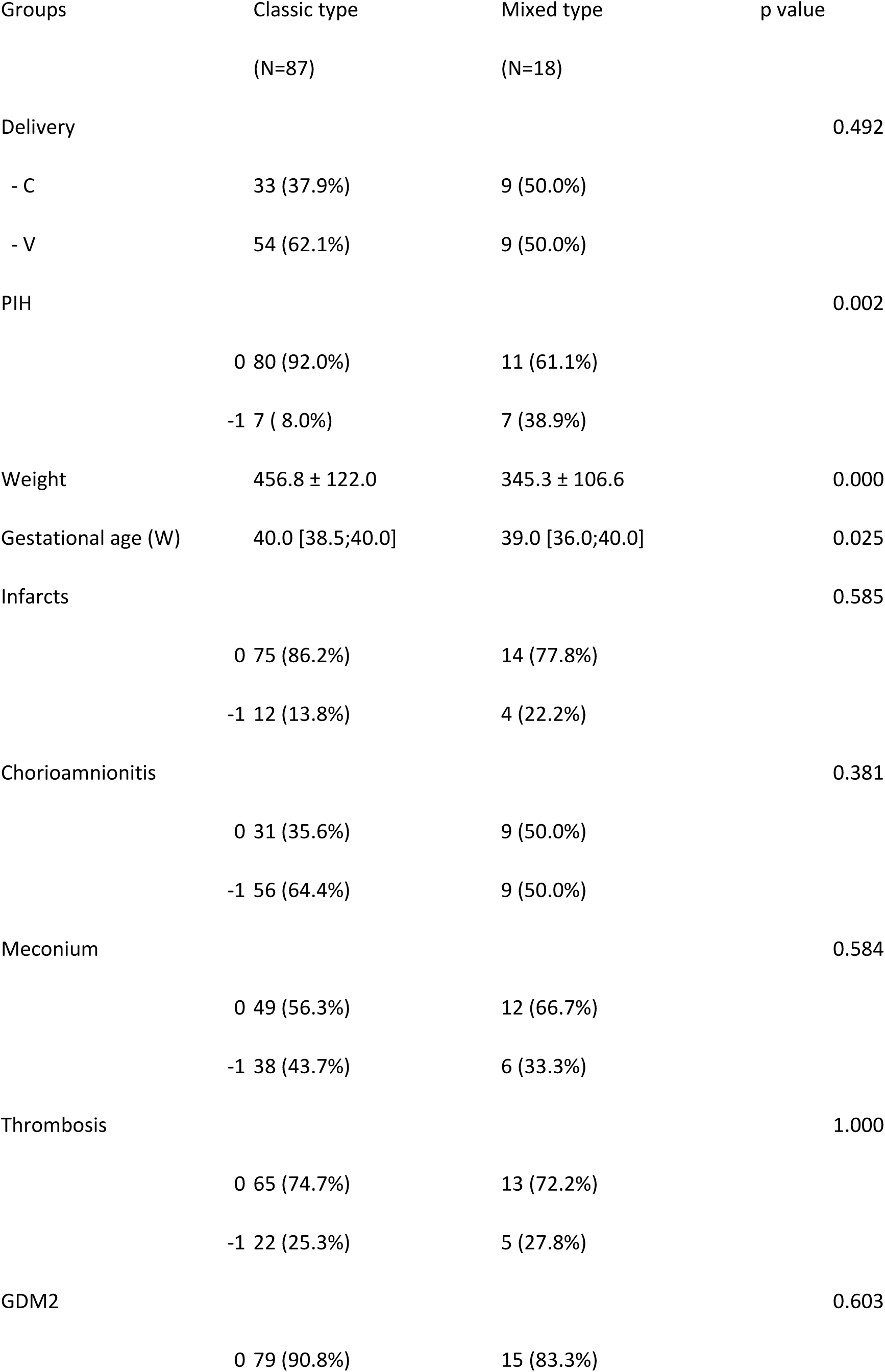
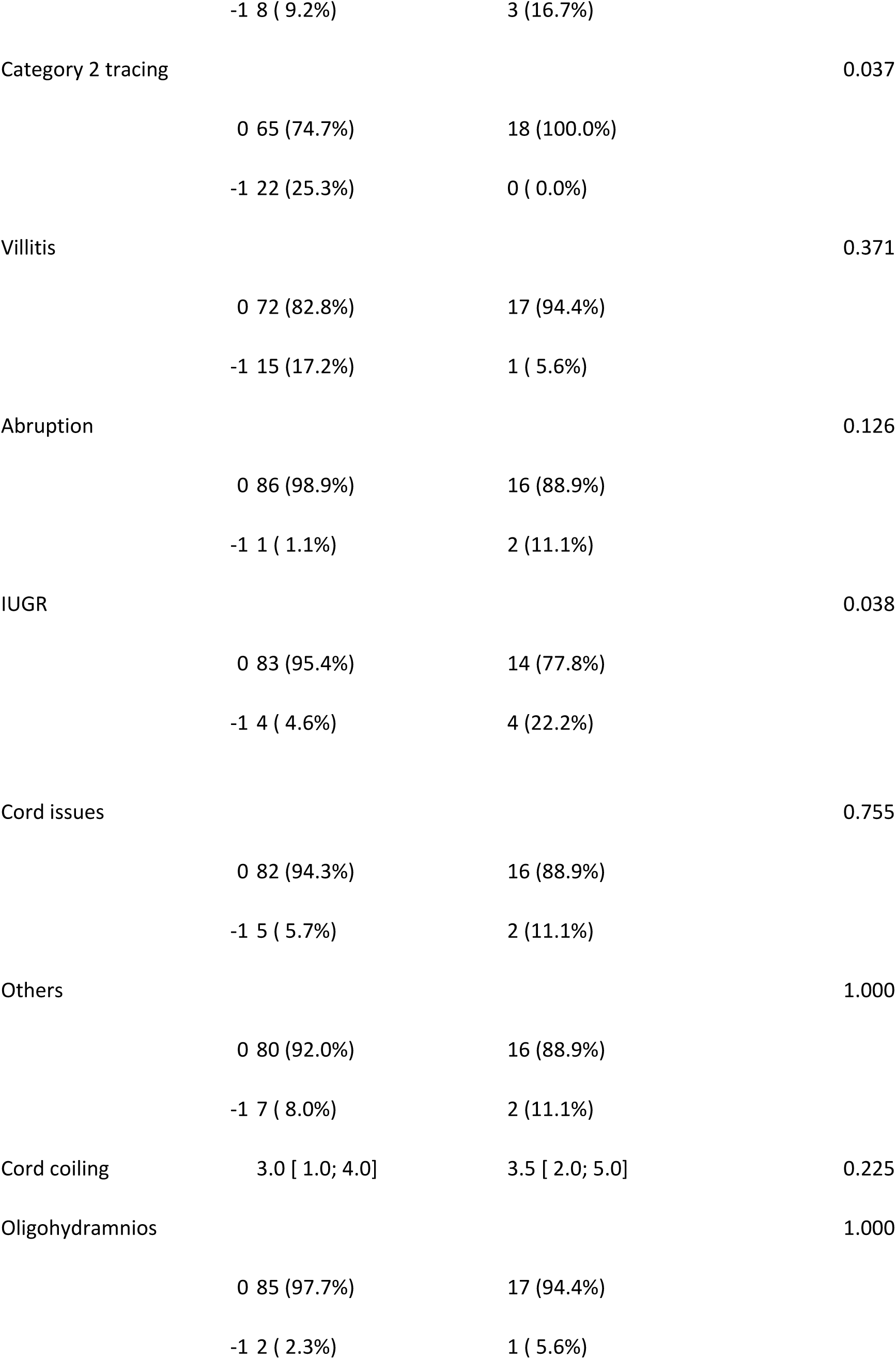
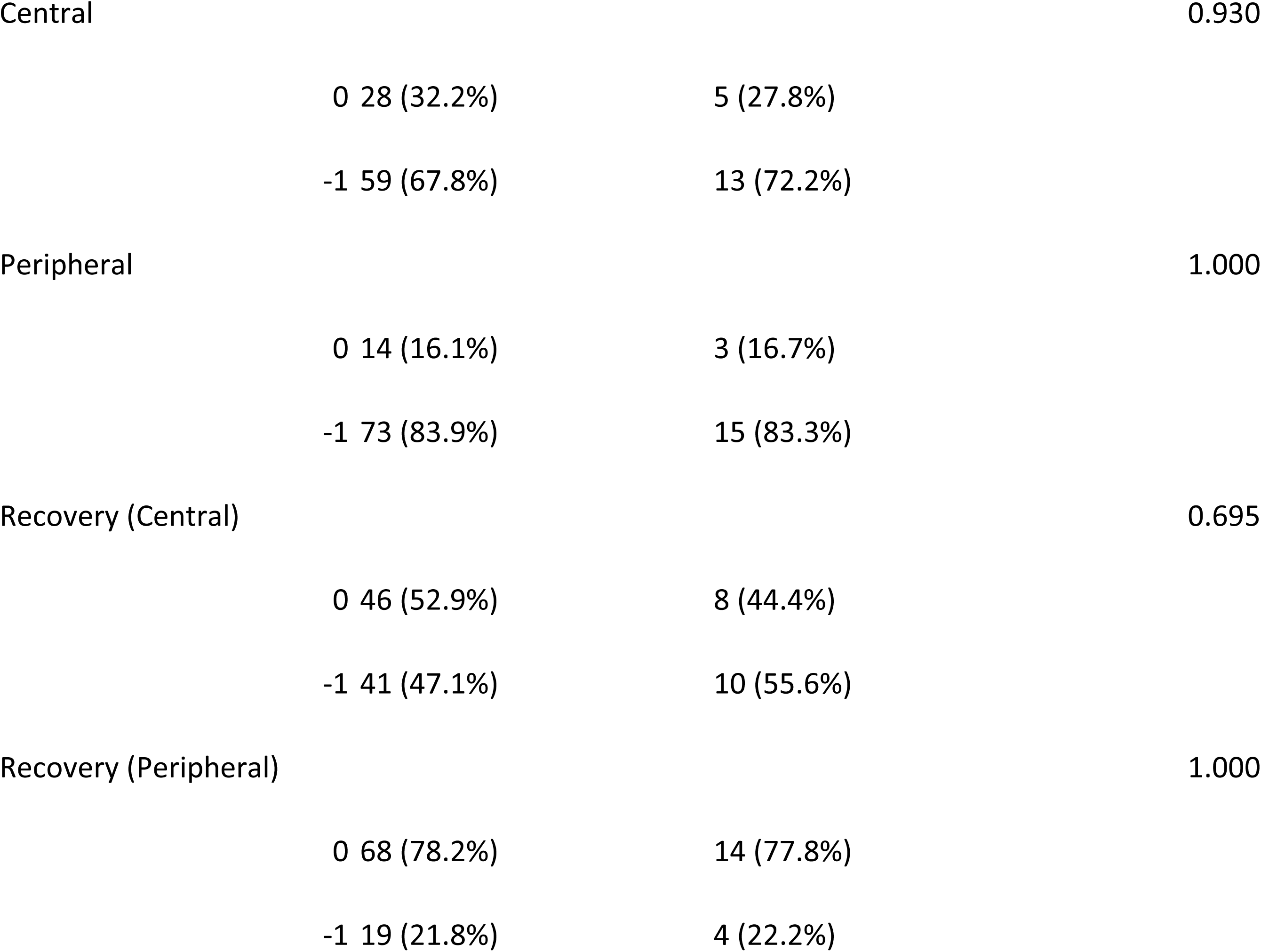
Baseline characteristics of 105 placentas with decidual vasculopathy and other clinical and pathologic findings within the placentas. 0= absence of the change, -1= presence of the change. Cord issues include abnormal insertions, knots, length, and cord vessels. Others include fetal anomalies, fetal vascular malperfusions, chorangiosis/chorangiomas, maternal gestational cholestasis, maternal autoimmune diseases, etc. Cord coiling refers to numbers of coils per 10 cm of cord.

**Table 2:** Frequency of decidual vasculopathy and vascular recovery (regeneration/restoration) in the central and the peripheral regions of the placentas with Fisher exact probability test (One tailed: p=0.0018, Two tailed: p=0.0024)

| Decidual Vasculopathy | Central | Peripheral | Recovery (C) | Recovery (P) | P value |
| --- | --- | --- | --- | --- | --- |
| Total | 72 | 88 | 51 | 23 | p=0.00049 |
| Classic | 59 | 73 | 41 | 19 | p=0.0018 |
| Mixed | 13 | 15 | 10 | 4 | P=0.113 |

**Figure 2.**
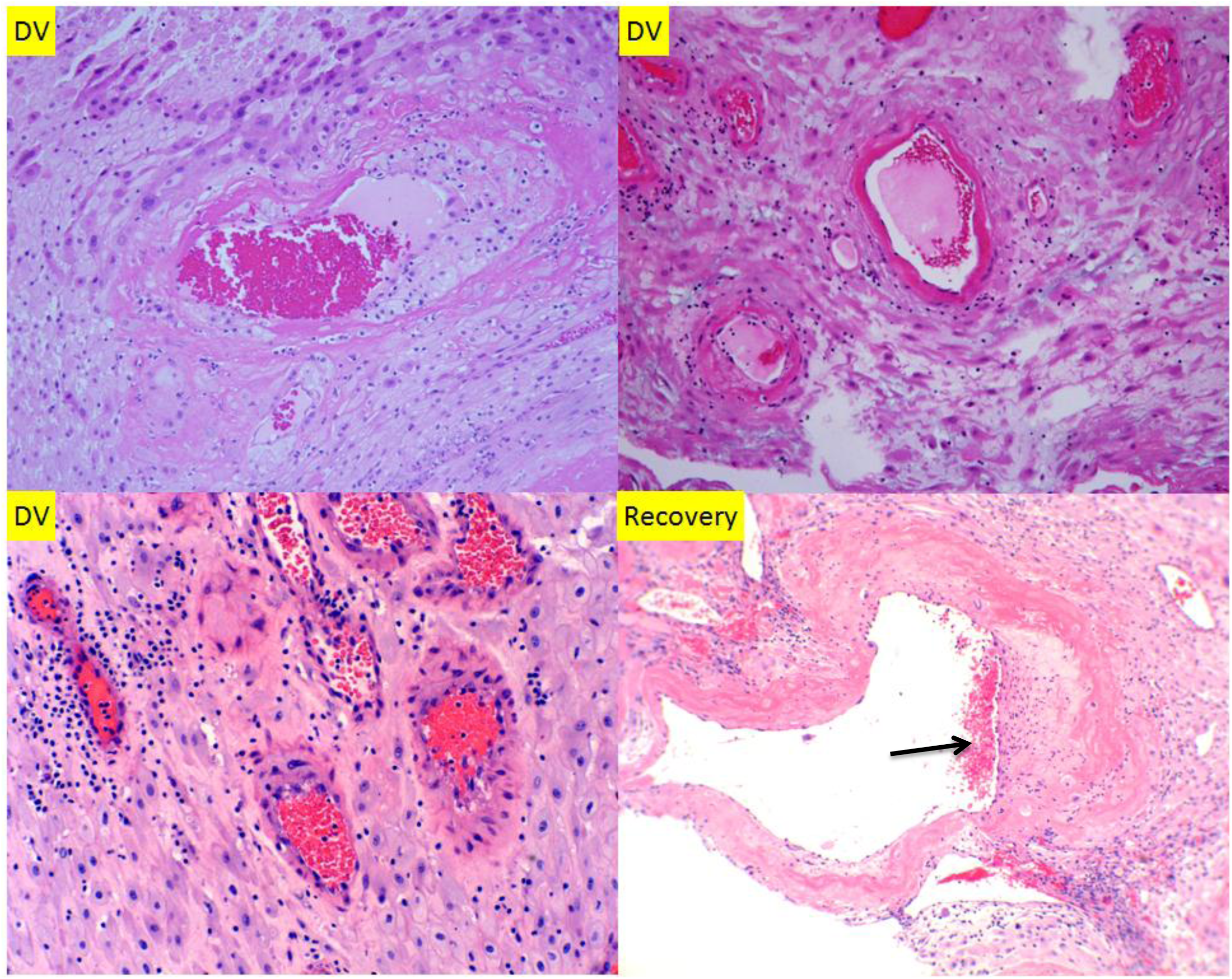
Classic decidual vasculopathy including acute atherosis and fibrinoid media necrosis with vascular regeneration. Mural arterial hypertrophy (bottom left) is trophoblasts-independent and therefore not included in the study. Black arrow points to “cushion” with smooth muscle cell regeneration. All sections were at 100 x magnification.

**Figure 3.**
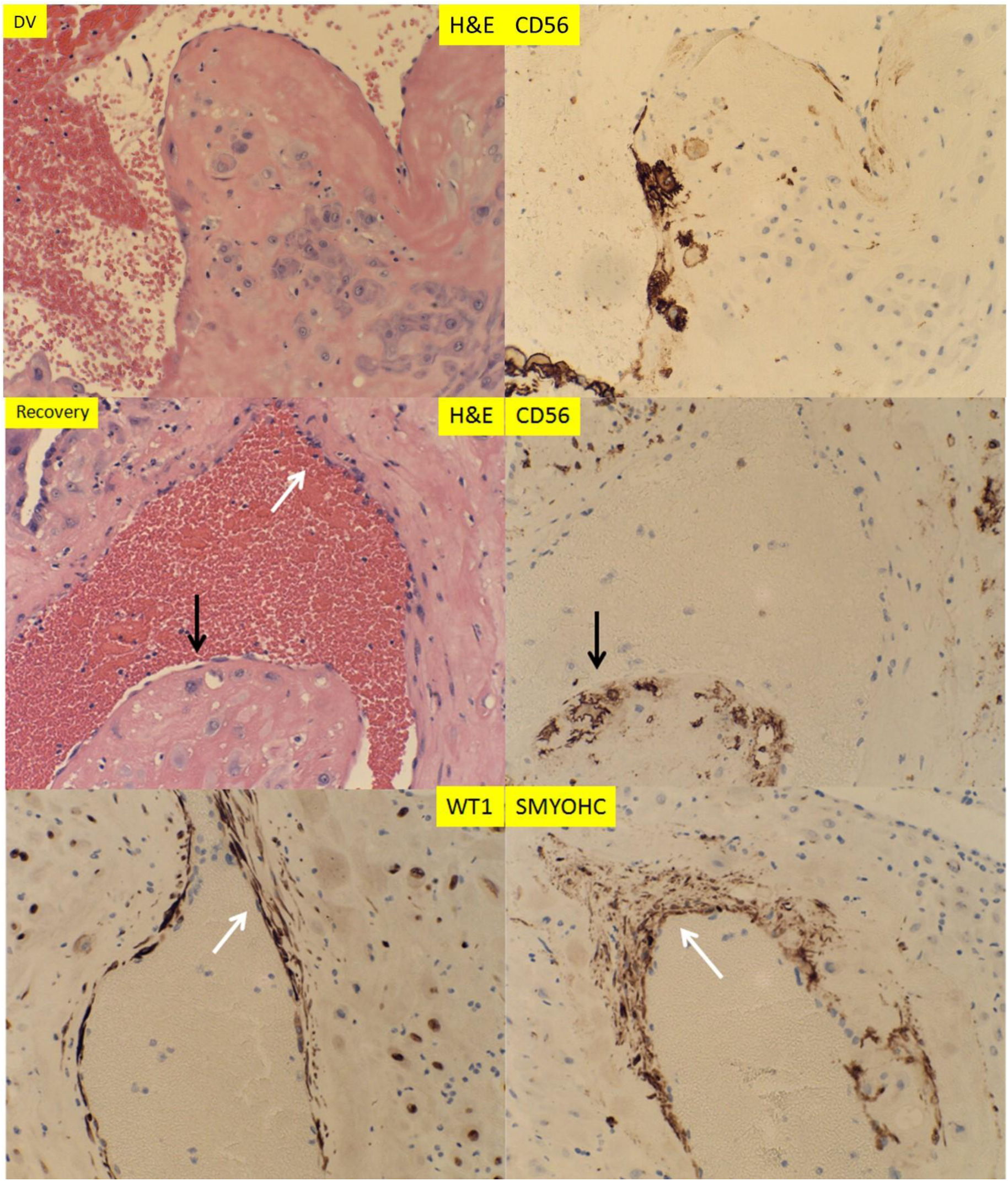
Representative sections of decidual vasculopathy and vascular regeneration / restoration with corresponding immunostaining patterns. The white arrows point to regenerating endothelial cells and smooth muscle cells and black arrows point to residual endovascular/intramural trophoblasts. All section were at 200 x magnification.

We examined one hysterectomy specimen containing the placenta at the 39 weeks with placenta increta within the previous C-section scar for vascular changes within the decidua and superficial myometrium (Figure 4). The patient didn’t have clinical history of preeclampsia or chronic hypertension. All of the sections of the myometrium and placental tissue were from the central areas. Decidual vasculopathy can be easily identified with various degrees of vascular regeneration /restoration within the decidual tissue (Figure 4 Top panel). The residual endovascular and intramural trophoblasts were detected by immunostaining for CD56, and the regenerating endothelial cells and smooth muscle cells by WT1 and SMYOHC (Figure 4). In contrast, the vascular changes within the superficial myometrium deep to the decidual tissue showed only regenerating arterial wall with endothelium and smooth muscle cells. No CD56-positive trophoblasts were detected in the myometrium section (Figure 4 bottom panel).

**Figure 4.**
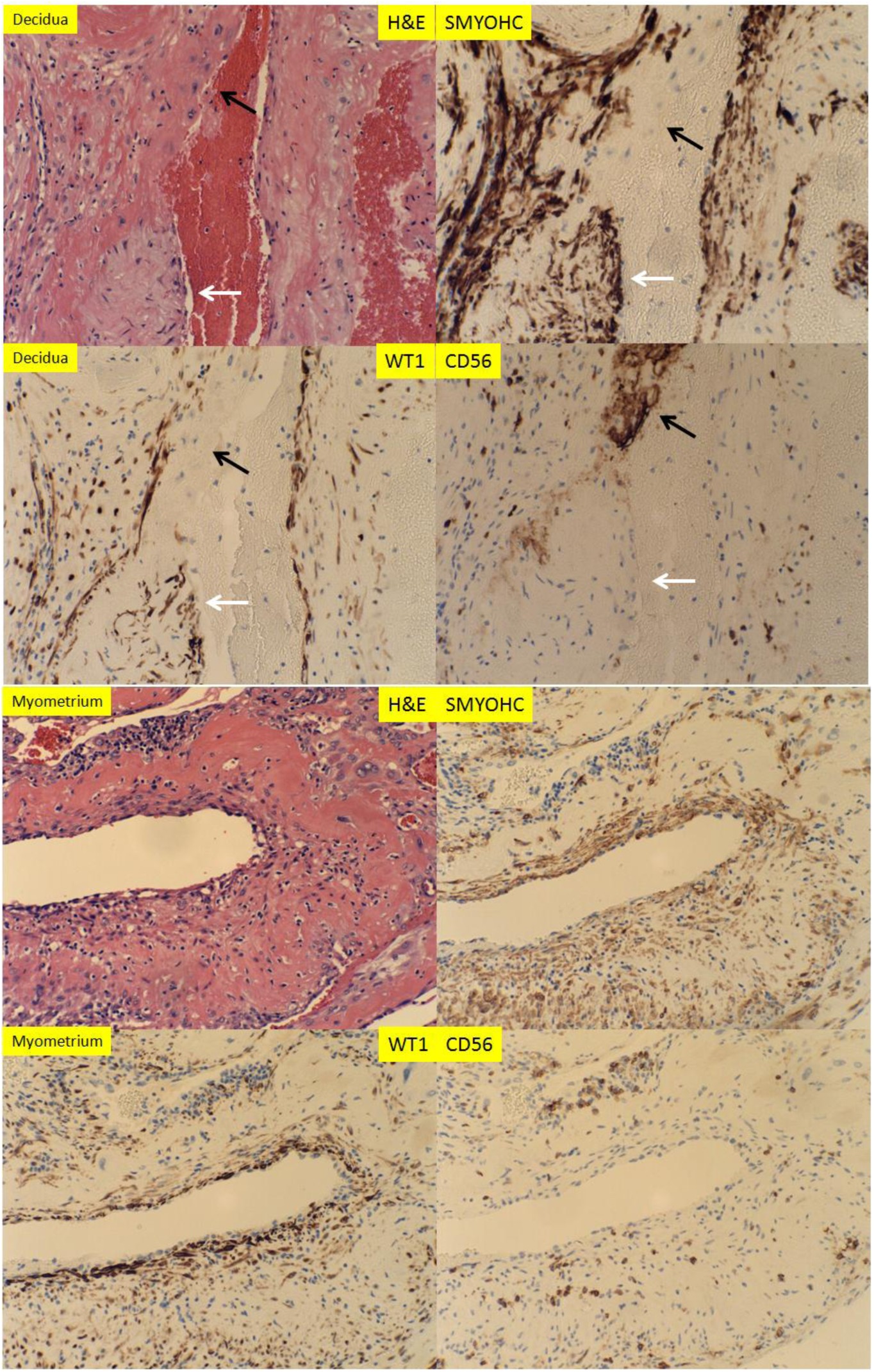
Histologic examination with immunostaining of vascular changes of spiral artery from hysterectomy specimen containing decidua and corresponding superficial myometrium. The white arrows point to regenerating endothelial cells and smooth muscle cells and the black arrows point to residual endovascular/intramural trophoblasts. All sections were at 200X magnification

## Discussion

In current study we attempted to evaluate the maternal surface for decidual vasculopathy in a spatial fashion to answer the question if there is a difference in morphologic changes of spiral artery that reflects the temporal vascular transformation. The study is based on the hypothesis that the central portion of the placenta is formed first with lateral expansion for continuous placental growth to term. There are few studies devoted to the lateral growth of placenta and spiral artery remodeling in a spatiotemporal fashion although much of the effort focused on the trophoblasts dependent remodeling of spiral artery in temporal fashion [11, 12]. Placenta is a transient organ, and maternal vascular transformation during pregnancy undergoes significant variations from the early trophoblastic plugging”, disappearance of smooth muscle wall and fibrinoid medial necrosis to the disappearance of endovascular trophoblasts and regeneration / restoration of endothelium and smooth muscle wall after delivery. Our previous study demonstrated that the intramural and endovascular trophoblasts persisted from early implantation to late pregnancy with formation of morphologic features of classic decidual vasculopathy including acute atherosis and medial fibrinoid necrosis [18, 22]. The link between decidual vasculopathy at late gestation to spiral artery remodeling at implantation raises the possibility of failure to die” of endovascular trophoblasts as a mechanism of pathogenesis of decidual vasculopathy. It also raises the questions if the death of endovascular trophoblasts is synchronously regulated through either maternal circulating factors (systemic) or specific interaction with the endothelial cells of the proximal artery (local) or both [22, 23]. It is frequently observed that the endovascular trophoblasts and the endothelium are mutually exclusive within the arterial lumen, but the intramural trophoblasts and endothelium can co-exist with overlying endothelium on the surface and underlying intramural trophoblasts within the arterial wall. This observation suggests that the endothelium regenerates /restores first after spiral artery remodeling of arterial wall. It is also consistent with the previous study that the endothelial progenitor cells give rise to the smooth muscle wall of the coronary artery [24, 25].

The formation of placental cotyledons from the center to the periphery is gradual and time-dependent. However, it is unclear if the formation of cotyledons is continuous, spanning the entire gestational age, although the placental weight increase occurs most in the third trimester. The connection of the cotyledons to maternal circulation through spiral artery remodeling from the center to the periphery should also theoretically be time-dependent. Our current data showed there is a statistically significant difference in regards to the frequency of decidual vasculopathy between the central portion and peripheral portion of the placenta, as well as the regenerating/restorative features of the spiral artery after decidual vasculopathy. The dynamic changes of decidual vascular transformation in a spatiotemporal fashion help interpretation of decidual vasculopathy during placental examination after delivery, as the placental examination only captures the snapshot of vascular changes of the spiral artery in specific time.

Pathogenesis of decidual vasculopathy is controversial and the factors affecting the cell death program of endovascular trophoblasts and relationship to regeneration/restoration of endothelium and smooth muscle wall are unclear at this time. Wilms tumor (WT1) gene and Wilms tumor associated protein (WTAP) gene expressions have been shown to be important for vascular and smooth muscle regeneration [21, 26, 27]. Smooth muscle regeneration in coronary arteries is dependent upon WT1 gene expression from the endothelial progenitor cells [24]. WT1 gene expression has been studied extensively in other cellular and animal models, and WT1 gene expression in endometrium during menstrual cycle and gestation was also attempted [20]. It is highly likely that quantitative measures are needed for WT1 gene expression in gestational vascular transformation to determine the roles of WT1 and WTAP genes in regeneration of endothelium and smooth muscles of the spiral artery locally, as a basal level of WT1 expression was detected within the endometrial stromal cells throughout the menstrual cycles and pregnancy [20]. It is also likely that unknown maternal circulating factors will influence the WT1 gene expression globally in a relatively synchronous manner. To search factors affecting WT1 gene expression within the regenerating endothelium and smooth muscles of spiral artery using in vitro models will likely provide useful information in pathogenesis of decidual vasculopathy and preeclampsia.

The controversy of vascular changes within the decidua and the superficial myometrium remains as placental bed biopsy predominantly studied the vessels within the myometrium [15, 28]. Myometrial vessels continue distally to become the decidual spiral artery. Vascular transformation during early pregnancy started at the decidual portion and progress proximally to the myometrial portion. Consequently regeneration during late pregnancy or after delivery started proximally. Our current study only captured the decidual vascular changes at late gestation, and spatiotemporal vascular transformation at early stages of pregnancy requires more studies.

## Conclusion

There is a statistically significant difference in both the incidence of decidual vasculopathy between the central and peripheral placenta and the regeneration / restoration of endothelium and smooth muscle of spiral artery due to the lateral growth pattern and corresponding spiral artery remodeling. Dynamic and spatiotemporal view of the spiral artery remodeling during pregnancy provides useful information in development of decidual vasculopathy at term.

## Acknowledgement

The author is thankful for Nina Yermolayeva, Cheryl Hernandez, Elizabeth Feygina and Missver Jocson for their technical assistance in preparation of placental specimens.

